# Germline mutation rate predicts cancer mortality across 37 vertebrate species

**DOI:** 10.1101/2023.08.13.553123

**Authors:** Stefania E. Kapsetaki, Zachary T. Compton, Walker Mellon, Orsolya Vincze, Mathieu Giraudeau, Tara M. Harrison, Lisa M. Abegglen, Amy M. Boddy, Carlo C. Maley, Joshua D. Schiffman

## Abstract

The explanation for why some species are more susceptible to cancer than others remains an area of intense investigation. Cancer evolves in part through the accumulation of mutations and, therefore, we hypothesized that germline mutation rates would be associated with cancer prevalence and mortality across species. We collected previously published data on germline mutation rate and cancer mortality data for 37 vertebrate species. Germline mutation rate was positively correlated with cancer mortality (*P* = 0.008). Why animals with increased germline mutation rates die more from cancer remains an open question, however they may benefit from close monitoring for tumors due to hereditary cancer predisposition syndromes. Early diagnoses of cancer in these species may increase their chances of treatment and overall survival.

## Introduction

In relatively fast changing environments, there may have been positive selection for an increase in germline mutation rate (1). This may have been accompanied by genetic hitchhiking and/or random genetic drift increasing the allele frequency of genes that protect organisms from developing cancer or perhaps even increase their chances of developing cancer. Cancer can evolve de novo in an individual, or it can arise in the background of an inherited allele that predisposes the individual to developing a cancer (2, 3). This is true for both humans and non-human animals (4–8). Germline mutations can be a causal or contributing factor in cancer development. For example, having a single *BRCA1* or *BRCA2* mutation in the germline increases the risk of breast cancer development in women to 60-80% (2). Other germline variants, such as a mutant mismatch repair allele, increase the somatic mutation rate which then leads to a dramatic increase in the risk of developing colorectal cancer (3). Although many non-hereditary cancers can be prevented by changes in an individual’s lifestyle, hereditary cancers are harder to prevent due to the “first hit” of cancer development in all their cells. Furthermore, a poor ability to prevent mutations in the germline may be associated with a poor ability to prevent somatic mutations, through faulty DNA synthesis fidelity or repair. This could indicate that species with a higher germline mutation rate have a higher vulnerability to cancer compared to species with a lower germline mutation rate, although this concept has not yet been tested.

Understanding the connection between inherited germline mutation rate and cancer mortality risk across different species may have a positive impact on the lives of animals through cancer screening programs and also offer new models for human genetic cancer predisposition syndromes. We focused on germline mutations across different species that were available in public datasets. We tested whether yearly germline mutation rates across 37 vertebrate species (including 23 mammalian species, 10 bird species, 3 reptilian species, and 1 species of Actinopterygii) (9) could explain the variability in cancer risk and death, using cancer mortality data from the Zoological Information Management System (ZIMS) software (https://species360.org/).

## Results & Discussion

We used previously published germline mutation rate data (68 species) (9) and tested for cancer mortality (37 matching species) using the ZIMS software. For a given species, cancer mortality is measured as the number of animals that died of cancer divided by the number of animals that died (including dying of cancer, but not of neonatal issues and parental neglect: see Methods). Merging the animals in both datasets, the common garter snake (*Thamnophis sirtalis*) had the highest germline mutation rate (1.27 × 10^−8^), and the house mouse (*Mus musculus*) had the highest cancer mortality (0.36). The snowy owl (*Bubo scandiacus*) had the lowest germline mutation rate (1.02 × 10^−10^), and the southern screamer (*Chauna torquata*), the White-faced saki monkey (*Pithecia pithecia*), and Darwin’s rhea (*Rhea pennata*) had the lowest cancer mortality (zero). We found that germline mutation rate was positively correlated with cancer mortality across 37 species with available cancer data (Fig. 1; PGLS: *P*-value: 0.0008, F-statistic = 7.69 on 1 and 35 DF; R^2^ = 0.13; lambda = 0.00006). The high germline mutation rate and cancer mortality in some species could be due to the older age of their fathers (10), since male germ cells undergo many more divisions than female germ cells.

**Figure 1.**
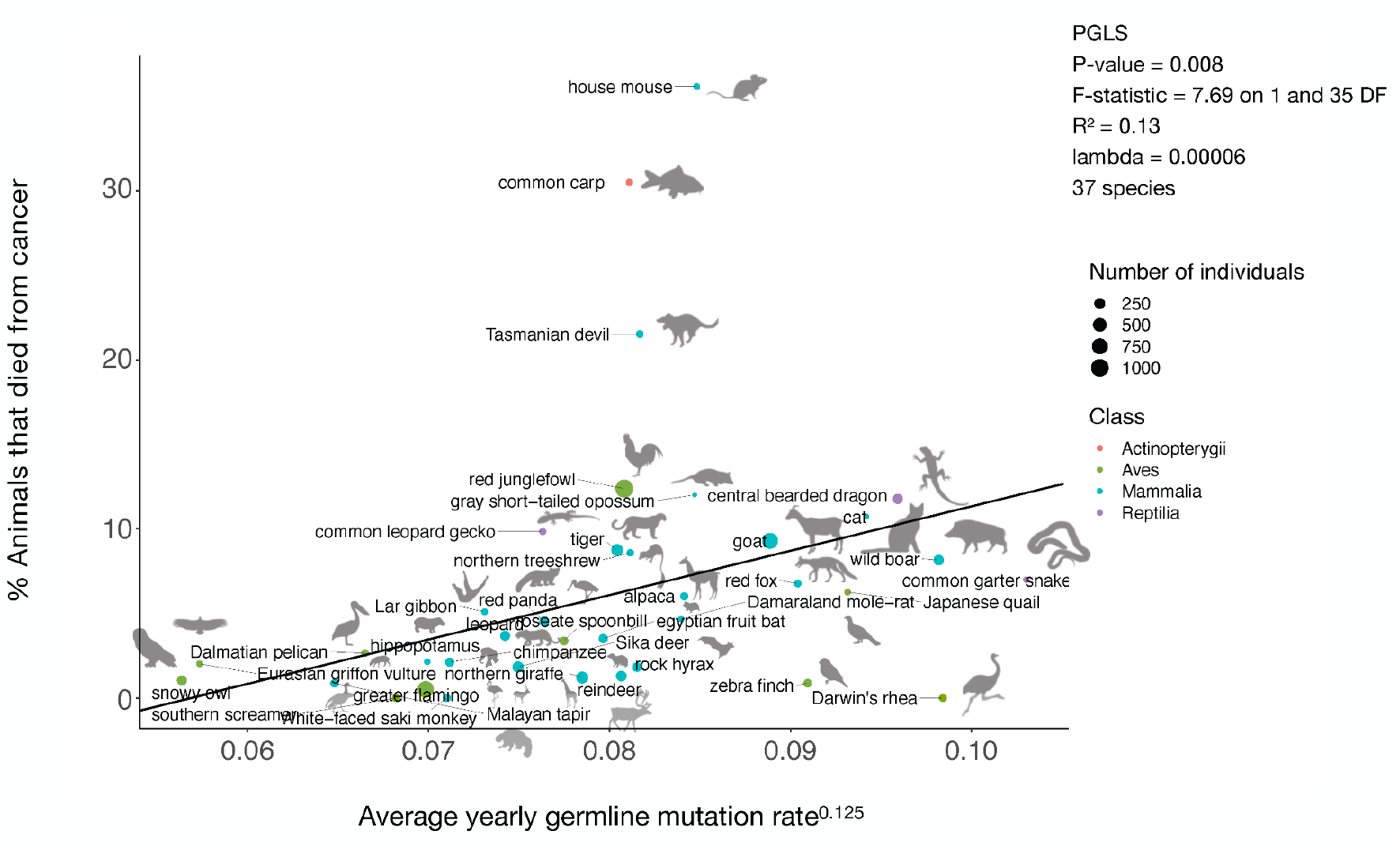
Average yearly germline mutation rate is positively correlated with the percentage of animals (among the total number of individuals per species examined at necropsy) that died from cancer. Each dot represents a species, and the size of the dot indicates the number of necropsies available for that species. The regression line is phylogenetically controlled (PGLS). Species’ images are from PhyloPic (https://www.phylopic.org/).

Cancer often appears after reproductive age in humans and other species which suggests strong selective forces in cancer defense mechanisms until later ages. Previous studies across vertebrates have found that cancer mortality risk is a trait under selection (4). Because cancer is often lethal, and can occur in animals that still have reproductive potential, we predict that the traits related to cancer suppression have likely evolved under natural selection as opposed to pure random genetic drift. We analyzed whether selection or genetic drift best explains patterns of germline mutation rate across the phylogeny of our 37 species. We found that the Ornstein-Uhlenbeck model of selection best fits our data (Fig. 2, AIC: OU = -1597.09; BM = -972.47; EB = -1148.26) showing that germline mutation rate is also a trait under stabilizing selection across the examined species. Previous analyses have identified trophic level (11), body size, and gestation length, (but not average adult lifespan) (4) as partial explanations for the variation in cancer prevalence across species, yet these factors only explain 1-31% of that variation. Our finding that the germline mutation rate explains 13% of the variation in cancer mortality across vertebrates (Fig. 1), suggests that it should be included in future efforts to understand cancer susceptibility across species. Although some connections (positive, negative, or none) between these variables are known (9, 12, 13), the exact associations between all variables still remain to be addressed.

**Figure 2.**
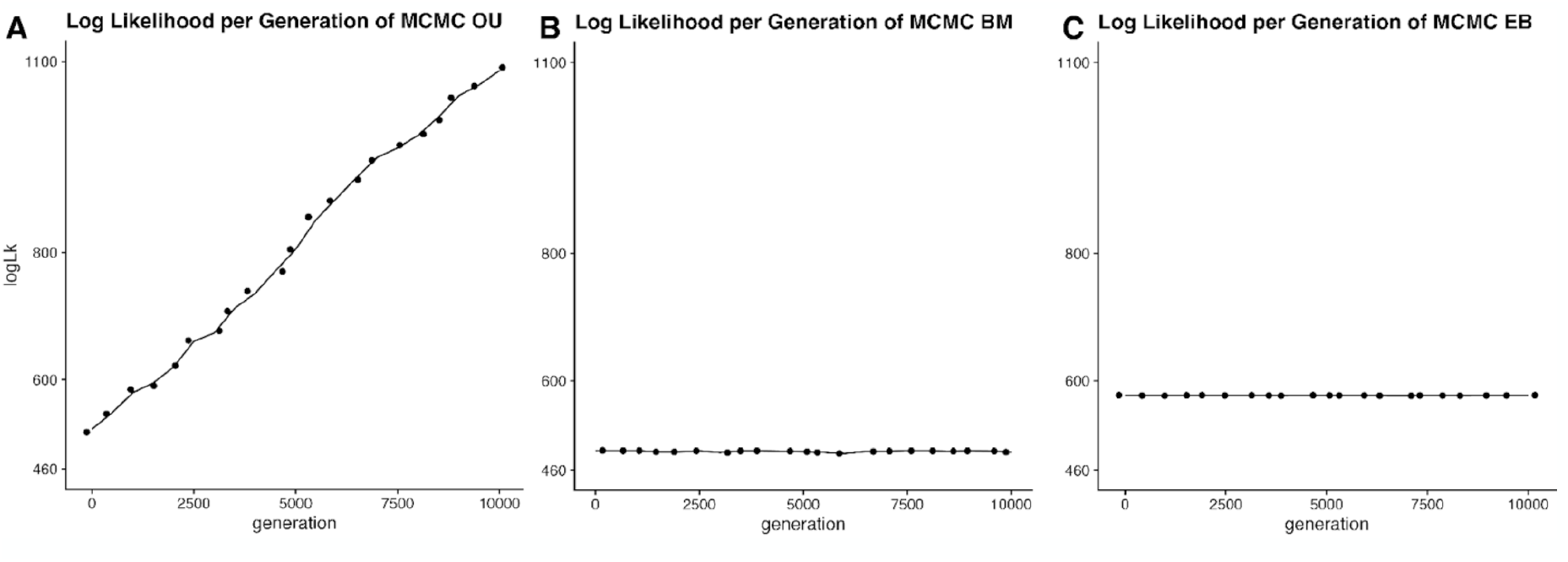
The Ornstein–Uhlenbeck (OU) evolutionary model (A), rather than the Brownian Model (BM) (B) or Early Burst (EB) model (C), best fits the species germline mutation rate data. Each point represents the reported log likelihood at each generation of the MCMC fitting algorithm. The OU model indicates that germline mutation rate is best described by a mode of evolution that reflects stabilization at an optimum value. The MCMC algorithm within the fitContinuousMCMC function generates samples of the parameter estimates from the posterior distribution. The likelihood is estimated at each generation of the algorithm to determine how well the parameter estimates fit the phylogenetic tree and the germline data.

The exact biological link between germline mutation rate and cancer mortality across vertebrates is unknown. There are many associations between germline mutations, such as mutations in *BRCA1/BRCA2* and *TP53*, and the occurrence of hereditary cancers in humans and dogs (14). Still, the causal relationship between germline mutation rate and cancer prevalence is unknown across vertebrates, i.e., whether an increased germline mutation rate is associated with an increased somatic mutation rate. Reproducing our results on an even larger number of species, with additional comparative oncology databases, and understanding the interactions between life history variables selecting for changes in germline mutation rate and cancer mortality, would help to answer our central question: what explains variation in cancer susceptibility across species? For now, the species we identified with an association between germline mutation rate and increased cancer mortality may harbor species-wide hereditary risk for cancer and thus may be under the risk of extinction. These species, in particular, may benefit from early cancer screening to diagnose tumors at an earlier and more treatable clinical stage of development.

## Methods

### Data Collection

Germline mutation rate data were from previously published literature (9). We estimated how cancer impacts species by using cancer mortality data. We estimated cancer mortality using the Mortality and Morbidity Module of ZIMS (https://species360.org/) software, from the periods of 1st of January 2011 to 13th of March 2023. To calculate cancer mortality for each species, we used the total number of individuals that died of cancer divided by the number of individuals that died of various factors (including cancer, but excluding the number of animals that died of neonatal issues and parental neglect, as an attempt to control for infant mortality that is likely to bias cancer risk estimates) as the denominator. We only used species with at least 20 records of mortality. If there was no report of neoplasia, we kept the numerator as zero. In both cases of the denominator and numerator, we only used the number of animals reported as having died of a single cause of death, not multiple causes of death.

### Models of evolution

We used three different Markov Chain Monte Carlo (MCMC) models of phenotype evolution (Ornstein–Uhlenbeck, Brownian, and Early Burst) to test which model was the best fit for the germline mutation data across the 37 species. To test for the best fit evolutionary model, we used the fitContinuousMCMC function from the *geiger* R package (15). The function utilizes species’ germline mutation data and our phylogeny to fit models using maximum likelihood. This version of the function, which utilizes MCMC, gives the test the ability to incorporate informative prior distributions for node values when the information is available.

### Statistical analysis

We performed the phylogenetic generalized least squares (PGLS) regression analysis in R version 4.0.5 using NCBI tree creator as described in previous studies (7, 11). The germline mutation rate data did not follow a normal distribution (Shapiro’s test), thus we transformed the data to the power of 0.125 (Tukey’s test) and then ‘centered’ this variable by subtracting it by its mean. We also weighted the PGLS by 1/(square root of the ZIMS denominator per species) to control for the variation in animal necropsies.

## Author contributions

A.M.B. and J.D.S. conceived the idea for this project. A.M.B. helped in guiding how to collect the cancer mortality data. S.E.K. collected the data, made figure 1, performed the regression analyses, and wrote the first draft. Z.T.C. and W.M. (under the guidance of Z.T.C. and C.C.M.) ran, compared, and analyzed the evolutionary models and made figure 2. A.M.B., C.C.M., T.M.H., and L.M.A. provided useful feedback throughout the project. All authors edited the final versions of the manuscript.

## Data and code availability

The data and code (https://github.com/zacharycompton/germlineMutation) will be available upon acceptance of the manuscript.

